# Spring ephemeral *Erythronium umbilicatum* may not be vulnerable to phenological mismatch with overstory trees

**DOI:** 10.1101/2025.05.14.652260

**Authors:** Melina Schopler, Anita Simha, Rebecca M. Dalton, Emma M. Wilson, Emmeline Redick, Elsa Youngsteadt, William K. Petry

**Affiliations:** Department of Plant and Microbial Biology, North Carolina State University, 100 Derieux Pl, Raleigh, NC 27607; Department of Biological Sciences, Louisiana State University, Life Sciences Bldg, Baton Rouge, LA 70808; Duke University Program in Ecology, Duke University, 2127 Campus Drive, Durham, NC, 27708; Department of Forestry and Environmental Resources, North Carolina State University, 2800 Faucette Dr, Raleigh, NC 27607; Department of Applied Ecology, North Carolina State University,100 Brooks Ave, Raleigh, NC 27607

**Keywords:** phenology, canopy leaf-out, shade experiment, historical data, temperature sensitivity, *Erythronium*

## Abstract

**Premise:** The defining life history strategy of spring ephemeral wildflowers is their avoidance of shading by trees during the brief, high-light period before canopy leaf-out. Studies suggest that spring ephemerals will experience increased light competition because canopy leaf-out is more sensitive to warming than is the phenology of spring ephemerals. However, it remains unclear how longer durations of shade will alter the population dynamics of spring ephemerals and whether all populations are at risk.

**Methods:** We experimentally shaded *Erythronium umbilicatum* for one to six additional weeks prior to canopy leaf-out to test for immediate and lagged effects of early shading on the timing of senescence and the probability of survival and flowering. To predict the potential for earlier shading, we combined long-term time series of spring air temperature, remotely-sensed tree leaf-out, and *E. umbilicatum* flowering phenology in North Carolina, USA.

**Key results:** Early shading did not alter *E. umbilicatum* until the following year, when more-shaded plants senesced later. We found no change in year-to-year survival, and a reduction in the probability of flowering only when plants experienced extremely early shading. Moreover, *E. umbilicatum* phenology was more sensitive than tree leaf-out to warming temperatures. Under climate warming, we project that *E. umbilicatum* is unlikely to experience shortened periods of high light.

**Conclusions:** Our findings show that a plant species’ defining life history strategy does not necessarily predict their sensitivity to phenological mismatches. This complicates, but also underscores the importance of identifying the most vulnerable species and directing our research efforts accordingly.

## INTRODUCTION

Plants often shift the timing of phenological events earlier in response to warmer spring temperatures, although some plant species respond more strongly to each degree of warming than others (Stuble et al., 2021). Species differences in phenological sensitivity to climate change can disrupt synchronized interactions or create new, historically absent interactions, a phenomenon known as phenological mismatch. Phenological mismatches, whether they exacerbate negative interactions or disrupt positive interactions, can have negative effects on fitness components and population trajectories. Spring ephemerals–perennial, understory herbs found in temperate, deciduous forests–exploit a short period of high light in the early spring when conditions are warm enough for aboveground growth but before the overstory canopy develops (Yancy et al., 2024). This shade avoidance strategy may be vulnerable to climatic changes that advance tree leaf-out faster than the onset of the growing season for spring ephemerals, resulting in unprecedented competition for light and a narrower window for growth and reproduction (Alecrim et al., 2022; Miller et al., 2023; reviewed in Lee et al., 2024). Understanding the multiyear effects of increased light competition on spring ephemerals and identifying which species are most likely to experience earlier canopy closure and where is crucial for directing our research and conservation efforts.

Since spring ephemerals exhibit a life history strategy evolved to escape light competition, and are only active above ground for a short period of time, any decrease in the duration of their high-light window would reduce their carbon budgets (Heberling et al., 2019) and should have an effect on their phenology, resource allocation, or fitness components. In *Erythronium* species, for example, the effects of shade on the current growing season include reduced fruit and seed set, reduced plant growth above and below ground, and earlier senescence (Kim et al., 2015; Augspurger & Salk, 2017). Moreover, reduced carbon budgets can exacerbate the physiological tradeoffs between investment in above-ground structures in the same year and in below-ground energy storage, which may affect survival and reproduction in the following year. However, no studies have experimentally examined the effects of early canopy shade on the following year fitness components. Because spring ephemerals are long-lived perennials, it is important to examine multiyear effects of shade on fitness components to understand how populations might be affected.

Despite evidence that reduced high-light windows can have negative consequences for some spring ephemeral species, the geographic locations where spring ephemerals face the greatest risk to climate change are still uncertain since different datasets bear conflicting results. In one study that spanned most of the eastern half of the United States, researchers used tree leaf-out date and spring ephemeral first flowering date from herbarium specimens to predict that light availability windows will decrease for spring ephemeral communities since tree leaf-out is more sensitive to warming spring temperatures than is spring ephemeral flowering (Lee et al., 2022; Miller et al., 2023). In contrast, Alecrim et al. (2022) used community science data from the National Phenology Network repository and found that light windows for spring ephemerals are increasing in duration at higher latitudes in North America, but remain constant at lower latitudes. Despite contradictory evidence from recent studies regarding where in North America spring ephemerals are most susceptible to early tree leaf-out (discussion of divergent findings can be found in Lee et al. 2023), both suggest that the phenological response to climate change is species and location specific. For example, while members of the genus *Erythronium* are some of the earliest to emerge and more sensitive to spring temperatures than other ephemerals, they could be found in forests dominated by trees with strong sensitivity to warming (such as *Acer rubrum* and *Fagus grandifolia*) which would still put them at risk of phenological mismatch with the canopy (Miller et al., 2023). Studies using alternative sources of data are needed to corroborate predictions from these models, which will help clarify when and where spring ephemerals may be most at risk.

To shed light on species-specific responses to early shade and location-specific interactions, we use *Erythronium umbilicatum*, a common species found in the southeastern United States and subject of several historical studies in Durham, NC (Motten, 1982, 1986). Members of this genus have been used in previous shade experiments although the lagged effects of shortened light windows have yet to be explored (Kim et al., 2015; Augspurger & Salk, 2017). In this paper we aim to: 1) test how early shade affects the current year and following year phenology and fitness components using an artificial shade experiment and 2) to understand the extent of phenological mismatch between *E. umbilicatum* flowering and canopy leaf-out using a novel combination of long-term ecological data and satellite imagery.

## MATERIALS AND METHODS

### Study site and focal species

We conducted this study at two sites in the Duke Forest in Durham, North Carolina, USA – Korstian Division: Gate 24 (35.983790, -79.032756; hereafter G24) and the Oosting Natural Area (35.980770, -79.064930; hereafter ONA). These sites are uniquely suited for this study since researchers previously monitored the flowering phenology of spring ephemerals at the same sites in 1978-1980 and again in 2015-2017 (Motten, 1986; Dalton unpublished data). The dominant tree species at these sites include *Acer rubrum, Fagus grandifolia, Liriodendron tulipifera*, and *Liquidambar styraciflua*.

We characterized the effects of early shade on *Erythronium umbilicatum* subsp. *umbilicatum* Parks and Hardin (Liliaceae), a spring ephemeral species common to the Piedmont region of North Carolina, with a range that extends north to Maryland and West Virginia, and south to Florida. *Erythronium umbilicatum* can be found in moist to mesic hardwood forests in floodplains and along gentle slopes (Radford et al., 2010). The life cycle of *E. umbilicatum* comprises two distinct phases– underground root and shoot growth that takes place during the autumn and winter, and aboveground growth that begins at the end of winter and continues through early spring (Lapointe, 2001). At our study sites, leaves and shoots typically emerge in late January or early February. During the spring epigeous growth period, plants photosynthesize and direct energy toward storage in the corm (the undifferentiated energy storage organ) or towards sexual reproduction in the form of flowers and fruits. Unlike other yellow *Erythronium* species of the eastern United States, *E. umbilicatum* does not reproduce asexually through rhizomes (Parks & Hardin, 1963).

In any given year, most individuals are vegetative at our sites (e.g., in our two-year study about 85% of individuals did not flower in either year). When flowering does occur, it often happens immediately after leaf emergence. Populations in the Piedmont of North Carolina reach peak flowering in March and finish flowering by the beginning of April. Each flower remains open from five to ten days. Common pollinators include a variety of native bees, particularly andrenid bees, and *Apis mellifera* (Motten, 1983).

### Aim 1: Effects of experimental shade on senescence, survival, and reproductive success

#### Experimental setup

We applied a gradient of early shade onset from six weeks ahead of natural canopy closure to no additional shading. We manipulated shade in one-week increments in replicate 1 m^2^ plots of naturally occurring patches of *E. umbilicatum*. For the shade treatment, we suspended shade cloth tents made of woven high density polyethylene (winemana, China) over treated plots. The cloths were designed to block >90% of solar radiation, mimicking full canopy leaf-out (>95% reduction in light levels; Augspurger et al., 2005). We oriented the open ends of the shade cloth tents to face north and south to maximize the shaded plot area over the course of the day. We also deployed a sham tent made of loosely-woven, light-transmitting nylon to control for potential fabric effects (e.g., on herbivores, pollinators, or precipitation; Jevrench, China; Appendix S1). We removed the sham and shade cloths after canopy leaf-out naturally occurred in 2023.

At the outset of the experiment in 2023, we identified all study plots and randomly assigned each to a treatment (uncovered true control, one to six weeks of sham control, or one to six weeks of shade). Beginning in the second week of February, we deployed five replicate shade cloths and one replicate sham each week over the course of six weeks, until the end of March. Each week, the shade and sham treatments were split between site G24 and ONA. We alternated which site received the extra shade cloth and which site received the sham plot. In total, we deployed up to five replicate plots for each of the six shade durations, one plot for each sham duration, and 12 replicate controls for a total of 45 plots across both sites (Appendix S2). Due to time constraints in the middle of the season, we switched to deploying treatments only at site ONA. In addition, we only deployed three shade replicates and no sham the week of March 20.

We tagged all *E. umbilicatum* individuals present in each plot (ranging from 4 - 30 individuals). In 2023, we visited the plots twice per week between February 20 and April 19, and recorded the date of senescence for each tagged plant. We considered a plant senescent if ≥ 50% of the total leaf area was yellow, if only the bare structural remains of the leaf were found (defined by Muller, 1978), or if the leaf was completely missing after signs of yellowing on prior observation days. In the first year of the study we only focused on the timing of senescence since some plants had already started to flower before we set up the plots. In 2024, we revisited the same plots twice per week, from the end of January through the end of April. We recorded if and when the tagged plants emerged, flowered, fruited, and senesced.

To evaluate the intended and unintended abiotic effects of the shade cloth, we measured the soil moisture as volumetric water content (VWC) in the southwest corner of a subset of plots at ONA on a fair-weather day in early March 2023 (Campbell Scientific, HydroSense II Handheld Soil Moisture Sensor; n_control_ = 24, n_shade_ = 9, n_sham_ = 2). We monitored light intensity (Lux) and soil surface temperature by deploying a pair of HOBO Pendant data loggers (Onset, HOBO UA-002-64 Pendant Light and Temperature Data Logger) in four shade plots and one sham plot at ONA. We placed the Pendants on top of the soil and leaf litter, with the light sensors facing upwards. Each pair consisted of a Pendant in the middle of the plot and one southwest of the plot, just outside the covered area. The Pendants recorded data every 10 minutes from March 28 - April 4, 2023.

### Analysis

#### 1. Shade calculations

For the first year of our study in 2023, when shade cloths were deployed on different dates, we calculated days of shade on a rolling basis. If the phenological observation was made before or on the day the treatment was deployed in a plot, the number of days of shade (or sham) for that observation was zero. If the observation was made after the treatment was deployed but before leaf-out, then the number of days of shade (or sham) for that observation was the number of days since the treatment was deployed. If the observation was collected after leaf-out in 2023 (day of year: DOY_leaf-out_ = 95 in 2023), then the number of extra days of shade (or sham) for that observation was calculated as the number of days the treatment preceded natural shade (*i*.*e*. 95 - DOY_treatment_). For phenological and fitness component measurements in 2024, we used the number of days the treatment preceded natural shade in 2023. In both years, all true control plots had zero days of extra shade.

#### 2. Environmental impacts of shade cloth

To quantify the impact of the shade and sham cloth treatments on light availability and air temperature, we compared daytime light and temperature (from 10:00 to 16:45) and nighttime temperature (19:00 to 04:45) of the paired pendants (n_shade_ = 4, n_sham_ = 1). We analyzed the data using two-way analysis of variance (ANOVAs) with a random intercept of plot and a first-order autoregressive variance structure to account for temporal autocorrelation of the residuals. We compared soil moisture values measured in all plots on March 6, 2023 using a one-way ANOVA.

#### 3. Phenology, survival, and fitness components

To quantify the effect of shade on the timing of senescence in 2023 and 2024, we fit binomial generalized linear mixed models (GLMMs) with a logit link and binomial error distribution in which the probability that an individual plant had senesced was predicted as a function of observation DOY, days of extra shade in 2023, and the interaction between them. Since each plot was assessed on multiple days, we included a random intercept of plot nested within site. We then performed a Type III Wald chi-square test to test for a significant interaction effect between observation DOY and number of days of extra shade.

To quantify the effect of shade in 2023 on the probability of survival, flowering, and fruiting given flowering in 2024, we fit binomial GLMMs with a logit link and binomial error distribution in which the probability of an individual emerging, flowering, or fruiting in 2024 was predicted as a function of days of extra shade in 2023. We included a random intercept of plot nested within site in our models.

To assess the impact of the sham cloth compared to ambient conditions, we ran the same analyses as above on the sham plot data and used days of extra sham (rather than days of extra shade) as the predictor.

### Aim 2: Estimate the rates at which spring ephemeral flowering and tree leaf-out phenology are changing over time

#### Spring ephemeral response to warming temperatures

We chose peak bloom as our phenological estimator for *Erythronium umbilicatum* because of its robustness to sample size and detection biases that impact other phenological indicators (Moussus et al., 2010). To model the day of peak bloom of *E. umbilicatum* each year as a function of temperature, we aggregated observations from datasets collected at our study sites between 1978-1980 by Motten (1982) and ourselves between 2015-2017 and 2023-2024. Motten visited G24 and ONA at two to three day intervals and recorded qualitatively when the *E. umblicatum* population was in full bloom at each site, defined as the range of dates between when floral abundance at each site began to increase and decrease most rapidly (Motten, 1982). For those years, we estimated the day of peak bloom at G24 and ONA as the median day within Motten’s reported full bloom range. In 2015, 2016, and 2017 R.D. visited 24 plots across sites G24 and ONA at two to three day intervals and recorded the number of flowering *E. umbilicatum* individuals per plot from January to the end of April. In 2023 and 2024 we repeated R.D.’s protocol to monitor flowering phenology in 45 plots across both sites. To estimate the day of year that peak flowering occurred between 2015 to 2024 we fit curves to the raw flowering data using Poisson regressions with a log link function and included the day of year squared as the predictor with a random intercept of plot (Appendix S3). We designated the maximum peak of the curves as the day of year that peak flowering occurred.

We then correlated peak bloom DOY from the three data sets with monthly, 4 km resolution PRISM temperature data for January through April from each observation year at each site (PRISM Climate Group, Oregon State University, 2022). Our study sites were located ca. 3 km apart in two adjacent grid cells. We associated each site’s peak bloom dates with temperatures from that site’s specific grid cell in each year. We performed model selection with Akaike Information Criterion, corrected (AICc) using the `dredge` function from the *‘*MuMIN’ package (Bartón, 2024) to identify the candidate model containing the mean monthly temperature or mean of consecutive monthly temperatures weighted by the number of days in each month, that best predicted peak flowering date (Appendix S4). We then took the best candidate model and used AICc to determine if including an interactive or additive effect of site improved the fit. We found that the mean air temperature of February through April, without an effect of site, best predicted peak bloom date of spring ephemerals. We used the fitted linear model (peak bloom DOY = *β*_0_ + *β*_1_ × temperature_Feb-April_; Figure 1, Appendix S5) to hindcast when peak bloom would occur over time from 1970 to 2024, given a mean February through April temperature obtained from PRISM (Figure 2).

**FIGURE 1.**
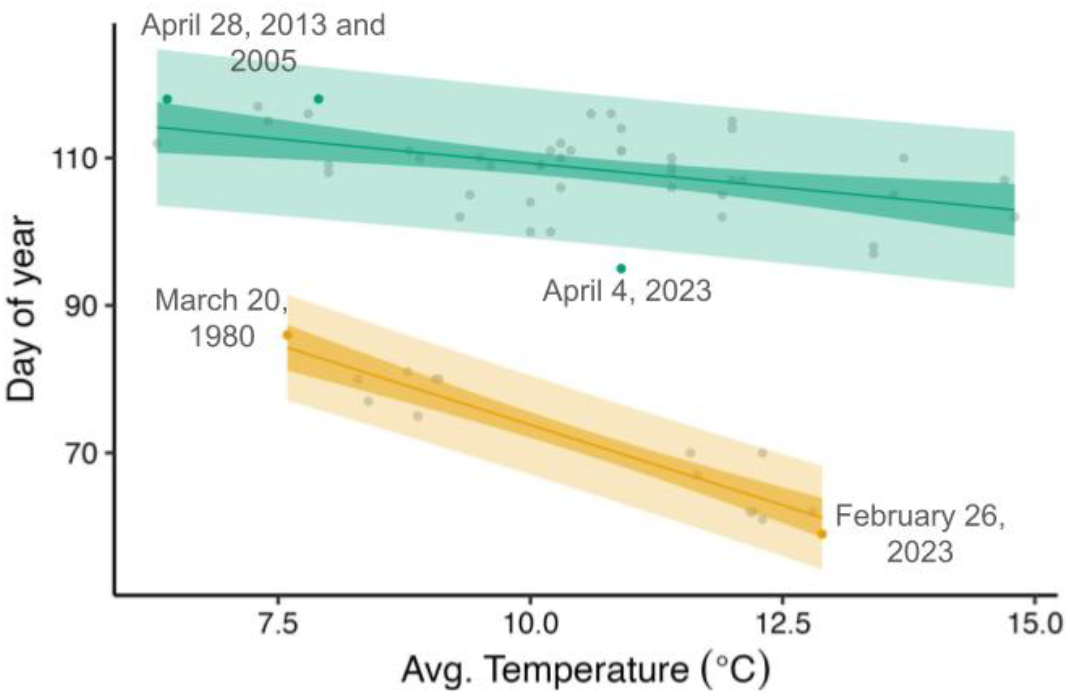
Best fit models to predict the day of year (DOY) that peak flowering and tree leaf-out will occur, given a mean spring temperature. Gold line (lower) represents *E. umbillicatum* peak flowering and green line (upper) represents tree leaf-out. The mean of February, March, and April temperatures were the strongest predictor of the timing of peak flowering. Mean March temperatures were the strongest predictor for the timing of tree leaf-out. Points represent raw data, lines represent fitted model with 95% confidence (dark ribbon) and prediction intervals (light ribbon). Colored points and text highlight the minimum and maximum observed day of year that the phenological events occurred. All temperature data were collected from the PRISM database.

**FIGURE 2.**
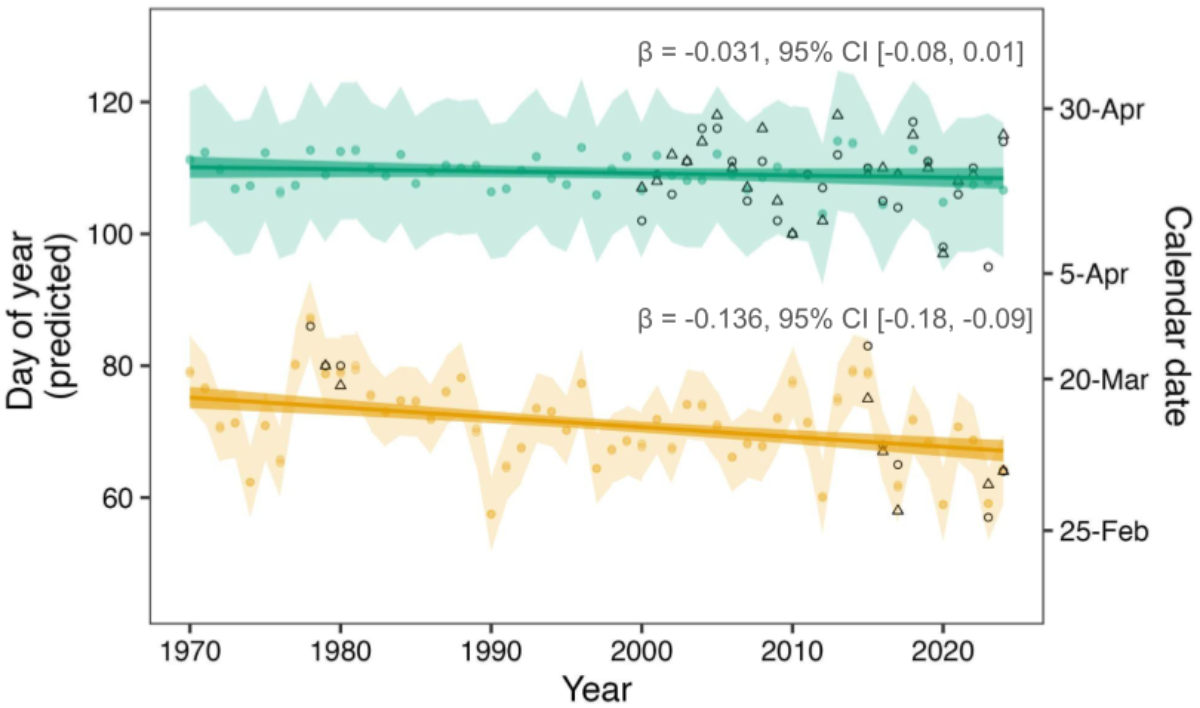
The flowering phenology of *E. umbilicatum* (gold) but not the timing of tree leaf-out (green) is predicted to shift earlier in the year 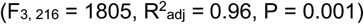. Linear regressions (shown in FIg. 1) were used to hindcast when a phenological event would have occurred in each year, given that year’s mean February-April temperature for flowering and March temperature for tree leaf-out. Colored points and jagged ribbons represent those predictions and 95% prediction intervals. For comparison to these predictions, open circles (G24) and triangles (ONA) represent direct observations of phenological events. These observations are the same raw data presented in Fig. 1, but were not involved directly in the linear fits shown here. Straight lines with 95% confidence intervals are outputs from the linear model that correlates year with predicted day of year of phenological event.

#### Tree leaf-out response to warming temperatures

We estimated the timing of tree leaf-out from 2000-2024 from the remotely-sensed MODIS Terra enhanced vegetation index (EVI) timeseries (MOD13Q1.061; Didan, 2021). The earliest available data were from the year 2000. The spatial resolution of the EVI data product is 250m and the temporal resolution is 16 days, which allowed us to differentiate our two study sites and extract six to eight EVI observations each year that spanned the leaf-out period (early February through the end of April). EVI is beneficial compared to other spectral indices because it retains its sensitivity in areas with a high leaf area index which is important for capturing more minute changes in leaf-out later in the season. When the tree canopy is fully closed, EVI is at its maximum. We first normalized the EVI data within each year and site to constrain data within zero and one (EVI_normalized_ = (EVI - EVI_min_) / (EVI_max_ - EVI_min_), where EVI_min_ is the minimum EVI value recorded at a site in a given year, and EVI_max_ is the maximum EVI value. We then used a quasibinomial GLM with a logit link to model normalized EVI as a function of year, site, DOY and their interactions (Appendix S6). Canopy leaf-out occurred rapidly; the EVI typically increased from <10% to >90% in about 56 days. We considered the inflection point of each line as the date of 50% green-up for each site and year, a proxy for tree leaf-out.

To find which months’ mean temperatures (or mean of consecutive monthly temperatures weighted by the number of days in each month) were the strongest predictors of the DOY that 50% green-up occurred, we obtained monthly mean temperatures from the PRISM database for each site for January through May from 2000 to 2024. We performed model selection with AICc using the `dredge` function from the *‘*MuMIN’ package (Bartón, 2024) in R (Appendix S4). We then took the best candidate model and used AICc to determine if including and interactive or additive effect of site improved the fit. We determined that mean March temperature, without an effect of site, was the strongest predictor of 50% green-up, and used the fitted model (50% green-up DOY = *β*_0_ + *β*_1_ × temperature_March_; Figure 1, Appendix S5) to hindcast the DOY of tree green-up for each year from 1970 to 2024, given a mean March temperature obtained from PRISM (Figure 2).

#### Comparing peak bloom and green-up response to warming air temperature

We used the two fitted linear models (50% green-up DOY = *β*_0_ + *β*_1_ × temperature_March_, peak bloom DOY = *β*_0_ + *β*_1_ × temperature_Feb-April_) to hindcast when the phenological events occurred each year, given a mean spring temperature from 1970 to 2024 (Figure 2). We then a fit linear model through those predicted points to estimate how those phenological events are trending over time (phenological event DOY = *β*_0_ + *β*_1_ × year × type; Figure 2). A greater sensitivity of *E. umbilicatum* to warming temperatures than that of tree leaf-out suggests a longer high light growing season as the climate warms and vice versa.

Previous studies commonly use a single, a-priori temperature variable to predict both flowering phenology and tree leaf-out (Alecrim et al., 2022; Miller et al., 2023), rather than using the strongest predictors for each phenological event. To investigate whether each method provides different results, we also modeled peak flowering time as a function of mean March temperatures (peak bloom DOY = *β*_0_ + *β*_1_ × temperature_March_; Appendix S5, S7). We used the fitted model to hindcast the DOY of peak flowering for each year from 1970 to 2024 and fit a linear model to those predicted points to estimate how the phenological events are trending over time (Appendix S8).

We performed all analyses in R 4.4.2 and RStudio 2024.04.1+748 (R Core Team, 2024; Posit team, 2024). We fit all generalized linear mixed models with the ‘lme4’ package (Bates et al., 2015). To analyze the Pendant data using a two-way ANOVA and a first-order autoregressive variance structure we used the ‘glmmTMB’ package (Brooks et al., 2017). We then used the ‘car’ package to perform Wald *X*^2^ tests to test for model significance (Fox and Weisburg, 2019). For post-hoc comparisons across treatments, we performed Tukey pairwise comparisons using the ‘emmeans*’* package (Lenth, 2024).

## RESULTS

### Aim 1: Effects of experimental shade on senescence, survival, and reproductive success

#### Environmental conditions of each treatment

Compared to ambient and sham conditions, the shade cloth significantly reduced visible light intensity by 80.5% (*χ*^2^ = 32.9, df = 1, P < 0.0001; Appendix S9) and daytime temperature by 6.6°C (*χ*^2^ = 12.6, df = 1, P < 0.0001; Fig. S10). Light levels and daytime temperature inside the sham plot were not significantly different to ambient conditions in a post-hoc Tukey test (Appendix S9, S10). The sham and shade cloths did not significantly affect nighttime temperatures (*χ*^2^ = 0.05, df = 1, P = 0.825) or soil moisture (F_2, 31_= 0.05, P = 0.974).

#### The effect of shade on timing of senescence and fitness components

The duration of additional shade did not affect the timing of plant senescence during the same growing season (*χ*^2^ = 1.40, df = 1, P = 0.237; Figure 3A, Appendix S11). Shade duration also did not affect survival in the following year (*χ*^2^ = 0.50, df = 1, P = 0.481). The effect of shade only became apparent later in the following growing season. Increasing shade duration in 2023 decreased the probability of flowering in 2024 (*χ*^2^ = 8.97, df = 1, P = 0.003; Figure 4; Appendix S12), although it did not affect fruiting given flowering (*χ*^2^ = 1.14, df = 1, P = 0.286). Shade in 2023 also delayed senescence in 2024 *(χ*^2^ = 5.06, df = 1, P = 0.025; Figure 3B, Appendix S11). Each week of additional shade in 2023 delayed senescence in the following year by approximately 0.6 days.

**FIGURE 3.**
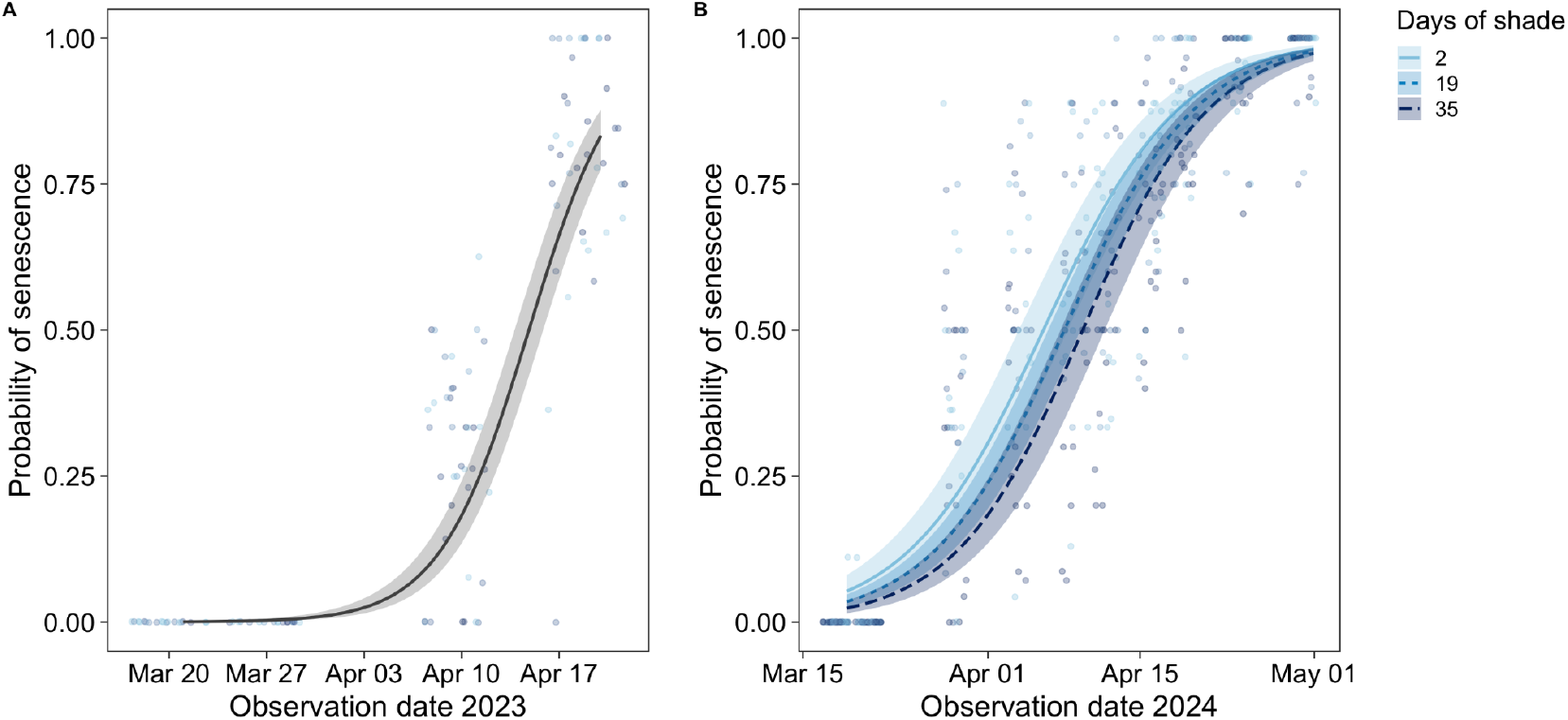
Experimental shade in 2023 (A) did not affect the timing of senescence in 2023 (P = 0.237) but (B) did affect the timing of senescence in 2024. Plants that received more days of shade in 2023 senesced later in 2024 (P = 0.025). The line in A depicts the logistic regression model fit across all shade treatments and the lines in B depict the mean number of days of extra shade (19 days) and ± one standard deviation number of days of extra shade (2 and 35 days). Points are raw data. Ribbons represent the upper and lower 95% CI.

**FIGURE 4.**
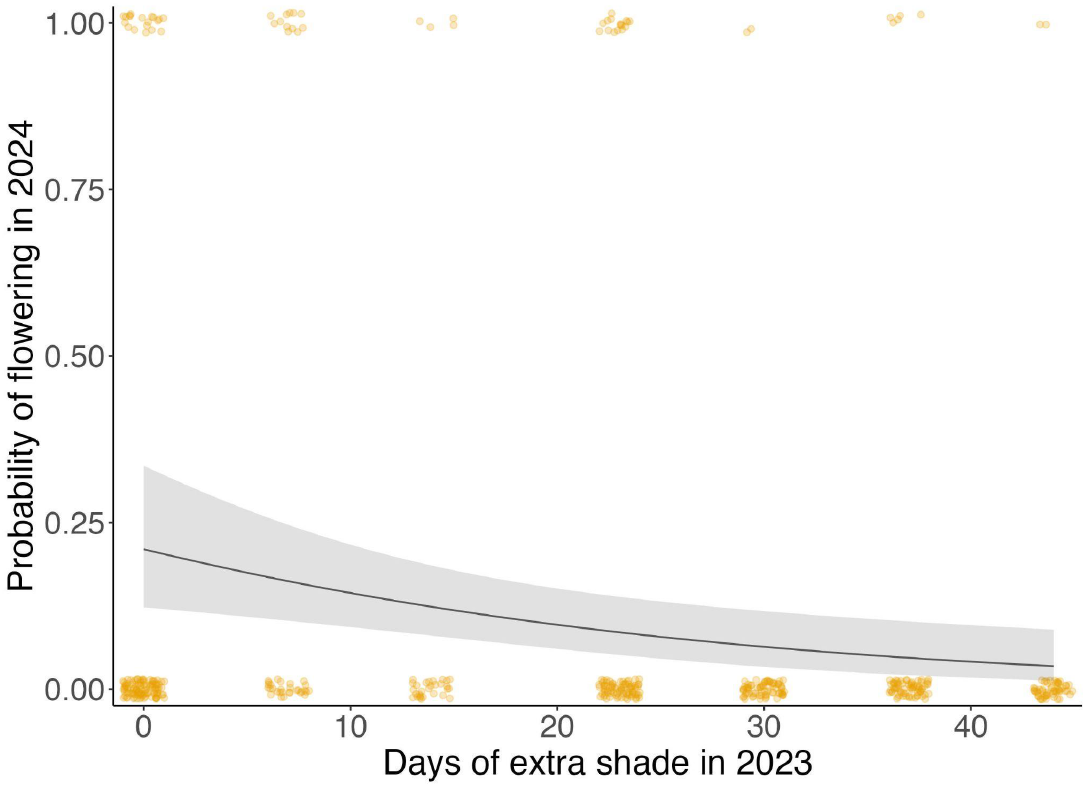
Longer experimental shade in 2023 decreased the probability of flowering in 2024 (P = 0.002). Points are individual plants (jittered to avoid overplotting), line is fitted model with 95% CI.

#### The effect of sham on timing of senescence and fitness components

The duration of sham cloth did not affect the timing of plant senescence during the same growing season (*χ*^2^ = 1.77, df = 1, P = 0.183). In the next growing season, 2023 shade had no lagged effects on 2024 survival (*χ*^2^ = 0.42, df = 1, P = 0.516), flowering (*χ*^2^ = 0.01, df = 1, P = 0.91), or senescence *(χ*^2^ = 0.80, df = 1, P = 0.371).

### Aim 2: The extent of phenological mismatch between *Erythronium umbilicatum* flowering and tree canopy leaf-out

The timing of *E. umbilicatum* flowering and tree leaf-out were best predicted by different temperature cues, and differed in their phenological sensitivity to their respective cues. For *E. umbilicatum*, the strongest predictor for peak flowering was the mean of February, March, and April temperatures (Appendix S4). For every °C of temperature increase, peak flowering occurred on average 4.35 (± 0.41) (mean ± S.E.) days earlier (*R*^*2*^ = 0.89; Figure 1, Appendix S5). The strongest predictor of 50% EVI green-up, our proxy for tree leaf-out, was the mean March air temperature (Appendix S4). For every °C of mean March temperature increase, 50% green-up occurred on average 1.3 (± 0.37) days earlier (*R*^*2*^ = 0.21; Figure 1, Appendix S5).

Using these temperature sensitivities to hindcast recent phenological change, we estimated that *E. umbilicatum* flowering has shifted earlier at a rate of 1.4 (± 0.5) days per decade, but the timing of tree leaf-out has not changed (*F*_*3, 216*_ = 1805, *p* = 0.001; Figure 2). Taken together, we estimate that the shade-free interval between *E. umbilicatum* peak flowering and tree leaf-out has increased by 1.4 days per decade.

When we modeled *E. umbilicatum* flowering as a function of mean March air temperature to investigate whether using a single temperature variable for both phenological events resulted in a different outcome, we found *E. umbilicatum* flowering was less sensitive to warming temperatures. For every °C of mean March temperature increase, peak flowering occurred on average only 2.72 (± 1.0) days earlier (*R*^*2*^ = 0.30; Appendix S5, S7). Moreover, using the mean March temperature sensitivity to hindcast recent change in flowering time reduced the rate of advancing flowering time to only .06 (± 0.05) days per decade (Appendix S8). There was also no significant difference between the timing of spring ephemeral flowering and tree leaf-out in response to warming temperatures over time (*i*.*e*. there was no significant interaction between type of phenological event and year; F_2, 216_ = 1577, p = 0.33).

## DISCUSSION

### Effect of shade on the timing of senescence

In contrast to what we would expect from carbon budgets and the results from previous shade experiments (Vezina & Grandtner, 1965; Kim et al., 2015; Augspurger & Salk, 2017), we found that early shade did not alter the timing of senescence in the year that the shade treatment was applied. We can conceive of two mechanisms for this surprising response. First, the shaded plants in our experiment may have acclimated to shade conditions, especially since our shade cloth only reduced light by 80%, when natural full canopy closure can reduce light by more than 90%. Relatedly, experimentally shaded leaves of *Erythronium japonicum* were larger and had a greater specific leaf area, which allowed them to continue to grow and maintain a carbon balance despite reduced light conditions (Kim et al., 2015). Another possible mechanism is that temperature might be a more influential than shade as a predictor of *E. umbilicatum* senescence. Lapointe (2001) found that senescence is triggered when carbohydrate reserves have been fulfilled, which occurs faster under warmer temperatures, regardless of the amount of shade.

Our shade treatment significantly delayed the timing of senescence in the following year. One possibility for this lagged effect of shade is that shaded plants did not store as much energy in their corm the year shading occurred (Muller, 1978), and extended leaf life span in the following year to compensate for reduced carbon acquisition and storage. The surprising lagged effect of shade on senescence in the following year emphasizes the need for additional multi-year studies to directly assess the mechanisms for delayed response to shade.

### Effect of shade on the demographic fitness components in the following year

Longer periods of shade did not alter *Erythronium* survival. Relatedly, previous work found that winter cold had a stronger effect on survival than did shade (Augsburger and Salk, 2017). While one year of extra shade did not reduce survival, repeated years of early canopy closure and decreased carbon acquisition might reduce individual survival over longer periods of time. Extra days of experimental shade did reduce the probability of flowering in the following year. A reduction in stored carbon under longer periods of shade could explain this result, since spring ephemerals rely on energy stored in the previous year for growth and flowering in the next (Routhier and Lapointe 2002). In control plots, the probability of flowering was 0.22. If light windows were to decrease by six days by 2080 (as projected for some spring ephemerals by Heberling et al., 2019), the probability of flowering would decrease by 20.5%. While a 20.5% reduction in the probability of flowering may seem extreme, the negative effects of early canopy shade may not be detrimental to *E. umbilicatum* populations since the intrinsic growth rate of long lived *Erythronium* species are least sensitive to fluctuations in the number of seedlings and more sensitive to relative growth rate and survival (Takada et al., 1998). To understand how early shade might impact spring ephemeral populations, future shade experiments should also study the effects of shade on following year demographic fitness components such as survival, growth, and reproduction.

### Assessing the risk of phenological mismatch

Although increased shade would likely reduce reproductive success in *E. umbilicatum*, the free-living populations in our area may not actually experience increased shade in the future. We found that *E. umbilicatum* is more sensitive to warming temperatures than are the canopy trees at our sites, thus the duration of the growing season might actually be increasing. On average, *E. umbilicatum* flowering is shifting 1.4 (± 0.5) days earlier per decade, whereas the trees are not leafing out any earlier. If this trend holds over time, *E. umbilicatum* in North Carolina could experience longer periods of high light in the spring than it has historically as the climate warms. Our findings are congruent with previous studies that found that *Erythronium* species are especially sensitive to warming temperatures and unlikely to experience decreasing light windows at lower latitudes in the short term as the climate warms (Alecrim et al., 2023).

While the results of our study suggest that *E. umbilicatum* may be resilient to the impacts of early shade, populations may still be vulnerable as the climate warms for several reasons. First, non-linear phenological responses to climate change can slow or even reverse species responses to warming (Iler et al., 2013; Pope et al., 2013), particularly with species that have a chilling requirement to break dormancy as do *Erythronium* spp. (Risser & Cottam, 1967; Yoshie & Yoshida, 1989). In the future, *Erythronium* species might not receive enough chilling degree days to break dormancy, resulting in delayed emergence and flowering as the climate warms. Second, emerging earlier each year could be dangerous for *E. umbilicatum* as late season frost events are becoming more common despite warmer average spring temperatures (Marino et al., 2011). More studies are needed on the effects of highly variable spring temperatures and late season frost events on the ability for spring ephemerals to shift their phenology. Finally, we investigated how *one* year of early shade affects phenology, survival, and reproduction in the next year. Under future climate conditions, spring ephemerals could experience multiple consecutive years of early canopy cover. More work needs to be done to understand the effects of consecutive years of early shade events.

One unique aspect of our study is that we used different temperature predictors in our models correlating mean spring temperatures with the phenological events, tree leaf-out and spring ephemeral peak flowering. We chose this approach since trees and spring ephemerals are known to be more sensitive to different climatic variables (Basler & Korner, 2014; Zohner et al., 2016; Jánosi et al., 2020). We suggest using the temperature predictors that are most relevant to each interacting group to best represent the mechanism by which phenological mismatch occurs. Phenological mismatch could arise when interacting species respond to the same cue but at different rates or if they respond to different cues. Studies that assume the former mechanism may under or overestimate phenological mismatch if the predictor variable is not a good fit. For example, with the present study, if we used mean March temperatures as the predictor of both phenological events, we would have estimated that the shade-free interval between *E. umbilicatum* peak flowering and tree leaf-out has not changed over time. Future studies could improve upon our approach by looking at how trees and spring ephemerals respond to different variables (such as accumulated growing degree days, soil temperature, chilling, precipitation, snowpack, photoperiod etc.) and the interactions between them.

In this study we also used the novel approach of combining long-term ecological data sets and satellite data to shed light on species-specific and location-specific interactions, which can be an effective method when tree and ephemeral data are not available from the same source. Both types of data sets are highly valuable and can be used in future studies to model the potential for phenological mismatch between trees and spring ephemerals globally. There is great potential for community science to accumulate similar data for other species and across large temporal and spatial scales (such as datasets from the USA National Phenology Network and Project Budburst). Going forward, it will be important to have standardized ways to collect data and record phenological events.

## CONCLUSIONS

It is critical to understand how threats such as climate change impact spring ephemerals, which are important for energy and nutrient cycling in temperate deciduous forests (Muller, 1978). Spring ephemerals are also important resources for many pollinator species, during a season when few floral resources are available (Dailey & Scott, 2006; Motten, 1986; Schemske et al., 1978).However, our understanding of which spring ephemeral species will be most sensitive to changing temperatures and how early shade will affect them is limited, which hinders our ability to protect and manage valuable ecosystems. Our results demonstrate that not all spring ephemerals will be sensitive to phenological mismatch with the canopy and highlight that a species’ defining life history traits do not necessarily predict vulnerability. Future studies are still needed to accurately predict the geographic and taxonomic variation in phenological responses to warming and the extent of ecosystem-level effects of changes in the spring-ephemeral community.

## Acknowledgments

The authors thank Director Sara Childs for supporting our research in the Duke Forest and Dr. Alexander F. Motten for familiarizing us with his old field sites and datasets. This work was supported by the Research Capacity Fund (HATCH), project award Nos. 7004646 and 7005517 from the U.S. Department of Agriculture’s National Institute of Food and Agriculture. Additional support was provided by the National Science Foundation Graduate Research Fellowship and Career-Life Balance Supplement to Melina Schopler under Grant No. DGE-2137100. Data collection for the years between 2015 and 2017 was supported by the Tom and Bruce Shinn Fund of the North Carolina Native Plant Society.

## Authorship Contributions

MS, WPK., EMW, and AS conceived of the research idea. MS, RMD, EMW, and ER collected and curated the data. MS performed the data analysis with input from EMW, EY, and WPK. MS wrote the manuscript with input from all authors. This work is not a product of the U.S. Government or the U.S. Environmental Protection Agency, and RMD is not doing this work in any governmental capacity. The views expressed are those of RMD only and do not necessarily represent those of the U.S. Government or the EPA.

## Data Availability Statement

The data from this study are openly available on Dryad at DOI: 10.5061/dryad.0k6djhbc9. Additional supporting information may be found online in the Supporting Information section at the end of the article.

## Appendix S1

Photographs of example sham shade control plot (white) and shade treatment plot (black) at site G24 located in the Korstian Division of the Duke Forest in Durham, North Carolina, USA.

## Appendix S2

Shade and sham control deployment schedule, locations, and replicate quantity in 2023. The number of days of extra shade that each plot received is the difference between the DOY that 50% green-up occurred in 2023 and the DOY that the treatment was deployed. In 2023 50% green-up occurred on DOY 95, which was April 5, 2023.

## Appendix S3

Poisson regression fit to counts of number of flowers per plot at each site in the specified years. Points represent the number of flowers in each plot (jittered to avoid overplotting). The maximum peak of each curve was used as an estimate of the day of year that peak flowering occurred.

## Appendix S4

List of months and consecutive combinations of months considered as predictors of each phenological event. Lowest AICc values bold, all other values are Δ AICc.

## Appendix S5

Output of linear models predicting the DOY that peak flowering and 50% green-up occurs as a function of mean spring temperatures. Mean February through April temperatures were the best predictor of the timing of peak flowing, whereas Mean March temperatures were the best predictors of the timing of canopy leaf-out. For comparison, we also estimated the DOY that peak flowering occurs as a function of mean March temperature.

## Appendix S6

Logistic regression fit to annual enhanced vegetation index (EVI) data from 2000 to 2024 to estimate when the canopy closed each year. Lines depict the mean fit of the model. Points depict raw EVI values (n = 6-8 observations per year). Each shade of green corresponds with a different year; lighter greens represent earlier years and darker greens represent later years. The 50% normalized EVI (subsequently called 50% green-up) was calculated as the inflection point of each line. G24 tended to leaf out slightly earlier than ONA in each year, although not statistically significant.

## Appendix S7

Linear models to predict the day of year (DOY) that peak flowering and tree leaf-out will occur, given a mean March temperature. Gold line (lower) represents peak flowering trend and green line (upper) represents tree leaf-out. Points represent raw data, lines represent fitted model with 95% confidence (dark ribbon) and prediction intervals (light ribbon). Colored points and text highlight the minimum and maximum observed day of year that the phenological events occurred. All temperature data were collected from the PRISM database.

## Appendix S8

The flowering phenology of *E. umbilicatum* (gold) but not the timing of tree leaf-out (green) is predicted to shift earlier in the year, although the slopes are not significantly different from each other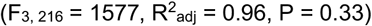. Linear regressions were used to predict when a phenological event would occur given a mean March temperature for both flowering and tree leaf-out. Colored points and jagged ribbons represent those predictions and 95% prediction intervals. Open circles (G24) and triangles (ONA) are the same raw data presented in Appendix S7, but were not involved directly in the linear fits shown here. Straight lines with 95% confidence intervals are outputs from linear model that correlates year with predicted day of year of phenological event.

## Appendix S9

The shade cloth significantly reduced the amount of light in the plots, while the sham light condition was not significantly different from ambient conditions (χ2 = 32.9, df = 1, P = <0.0001). The light gray points depict the raw data on the log scale. Boxplots summarize the data, error bars represent the estimated marginal means and 95% CI.

## Appendix S10

The shade cloth significantly decreased daytime temperature compared to ambient and sham conditions (χ2 = 12.59 df = 1, P = <0.0001). Light gray points are data, boxplots summarize the data, error bars represent the estimated marginal means and 95% CI.

## Appendix S11

The effects of observation day (center scaled), the number of days of extra shade, and the interaction between them in 2023 on the probability of senescence in 2023 and 2024. P-values significant at α = 0.05 are in bold font.

## Appendix S12

The effect of the number of days of extra shade in 2023 on the probability of flowering in 2024. P-values significant at α = 0.05 are in bold font.

