## Appendix S1 for "Spring ephemeral *Erythronium umbilicatum* may not be vulnerable to phenological mismatch with overstory trees"

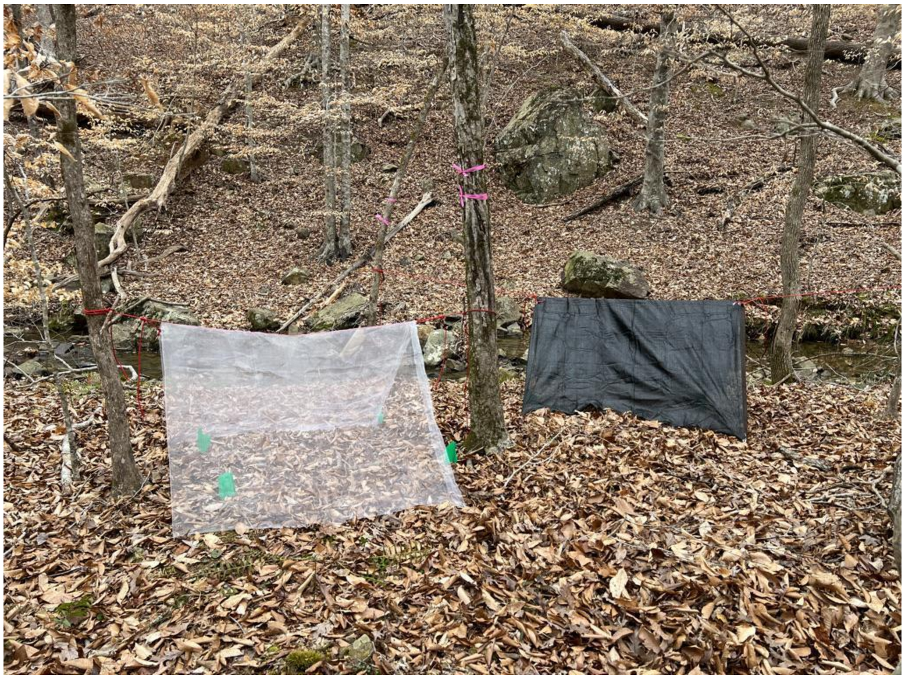

**Appendix S1.** Photographs of example sham shade control plot (white) and shade treatment plot (black) at site G24 located in the Korstian Division of the Duke Forest in Durham, North Carolina, USA.
