## Appendix S2 for "Spring ephemeral *Erythronium umbilicatum* may not be vulnerable to phenological mismatch with overstory trees"

**Appendix S2.** Shade and sham control deployment schedule, locations, and replicate quantity in 2023. The number of days of extra shade that each plot received is the difference between the DOY that 50% green-up occurred in 2023 and the DOY that the treatment was deployed. In 2023 50% green-up occurred on DOY 95, which was April 5, 2023.

| Treatment | Date deployed | DOY | Location(s) | Number of plots | Days of extra shade |
|---|---|---|---|---|---|
| control | NA | NA | 9 plots at G24; 3 plots at ONA | 12 | 0 |
| shade1 | 2-20-2023 | 51 | 3 plots at G24; 2 plots at ONA | 5 | 44 |
| sham1 | 2-20-2023 | 51 | G24 | 1 | |
| shade2 | 2-27-2023 | 58 | 2 plots at G24; 3 plots at ONA | 5 | 37 |
| sham2 | 2-27-2023 | 58 | ONA | 1 | |
| shade3 | 3-6-2023 | 65 | 3 plots at G24; 2 plots at ONA | 5 | 30 |
| sham3 | 3-6-2023 | 65 | G24 | 1 | |
| shade4 | 3-13-2023 | 72 | 2 plots at G24; 3 plots at ONA | 5 | 23 |
| sham4 | 3-13-2023 | 72 | ONA | 1 | |
| shade5 | 3-20-3023 | 79 | ONA | 3 | 16 |
| sham5 | none | NA | NA | 0 | |
| shade6 | 3-27-2023 | 86 | ONA | 5 | 9 |
| sham6 | 3-27-2023 | 86 | ONA | 1 | |
