## Appendix S3 for "Spring ephemeral *Erythronium umbilicatum* may not be vulnerable to phenological mismatch with overstory trees"

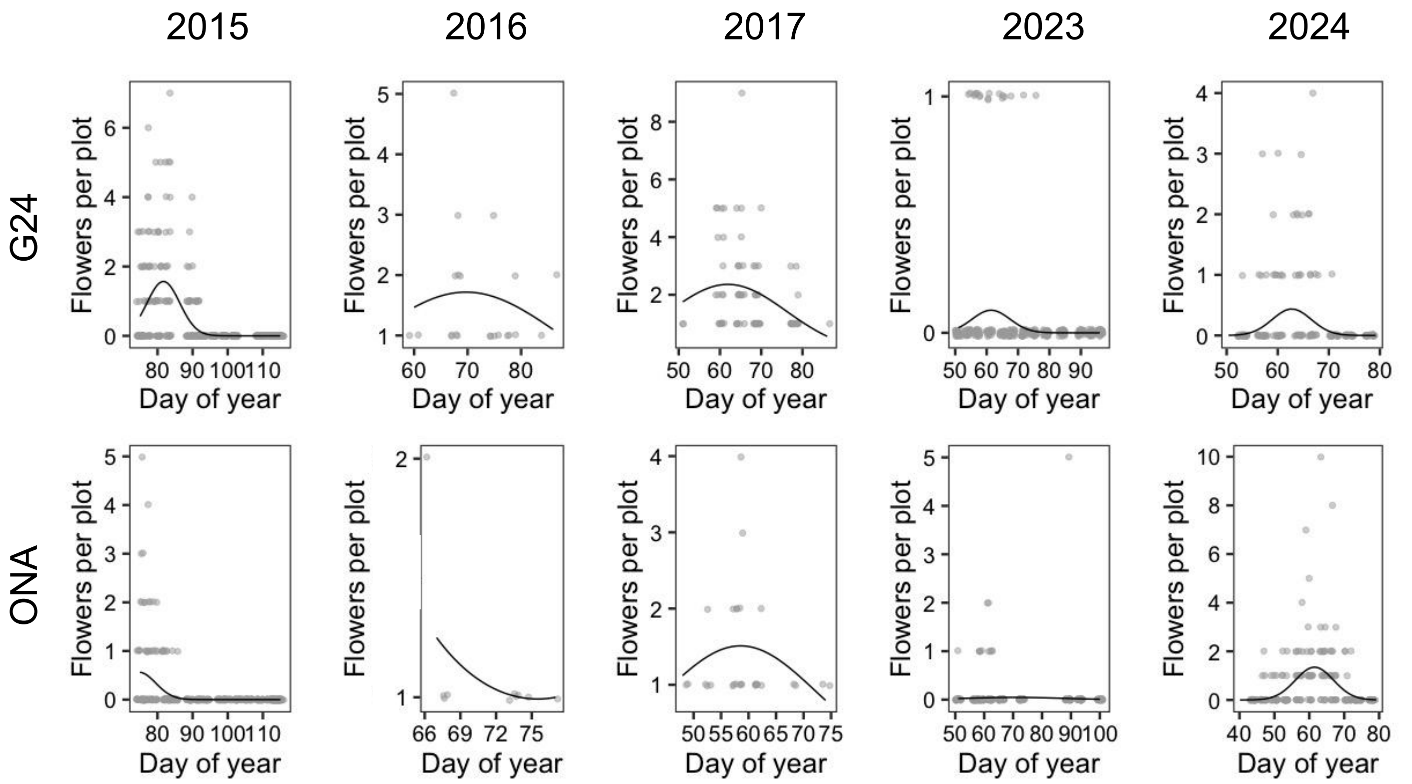


**Appendix S3.** Poisson regression fit to counts of number of flowers per plot at each site in the specified years. Points represent the number of flowers in each plot (jittered to avoid overplotting). The maximum peak of each curve was used as an estimate of the day of year that peak flowering occurred.
