## Appendix S4 for "Spring ephemeral *Erythronium umbilicatum* may not be vulnerable to phenological mismatch with overstory trees"

**Appendix S4.** List of months and consecutive combinations of months considered as predictors of each phenological event. Lowest AICc values bold, all other values are Δ AICc.

| **Peak Flowering** | | **50 % green-up** | |
|---|---|---|---|
| **Time period** | **AICc or Δ AICc** | **Time period** | **AICc or Δ AICc** |
| Feb-April | **81.51** | March | **282.77** |
| Jan-April | 1.02 | March-April | 0.26 |
| Jan-March | 4.39 | Jan-March | 5.12 |
| Feb-March | 5.76 | Jan-April | 5.50 |
| Feb | 6.10 | Feb-April | 6.03 |
| Jan-Feb | 10.34 | Feb-March | 6.08 |
| March-April | 16.95 | Jan | 9.28 |
| April | 20.53 | Jan-Feb | 10.78 |
| Jan | 21.57 | April | 11.54 |
| March | 27.41 | Feb | 11.68 |
