## Appendix S5 for "Spring ephemeral *Erythronium umbilicatum* may not be vulnerable to phenological mismatch with overstory trees"

**Appendix S5.** Output of linear models predicting the DOY that peak flowering and 50% green-up occurs as a function of mean spring temperatures. Mean February through April temperatures were the best predictor of the timing of peak flowing, whereas Mean March temperatures were the best predictors of the timing of canopy leaf-out. For comparison, we also estimated the DOY that peak flowering occurs as a function of mean March temperature.

| **Response** | **Predictor** | **Estimate** | **SE** | $R^2$ |
|---|---|---|---|---|
| DOY peak flowering | (Intercept) | 117.296 | 4.399 | |
| | Feb-April mean | -4.351 | 0.411 | 0.888 |
| | March mean | -2.72 | 1.02 | 0.304 |
| DOY 50% green-up | (Intercept) | 122 | 3.943 | |
| | March mean | -1.317 | 0.369 | 0.207 |
