## Appendix S6 for "Spring ephemeral *Erythronium umbilicatum* may not be vulnerable to phenological mismatch with overstory trees"

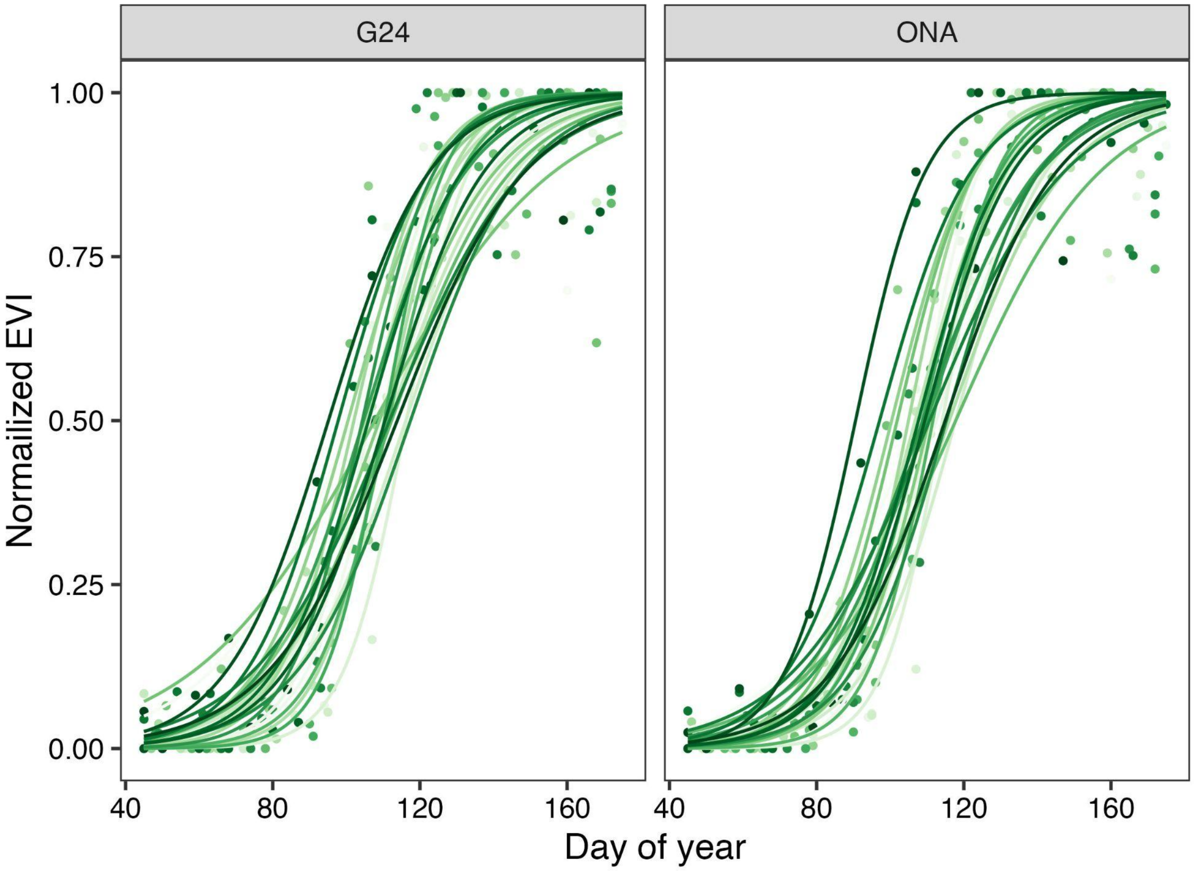


**Appendix S6.** Logistic regression fit to annual enhanced vegetation index (EVI) data from 2000 to 2024 to estimate when the canopy closed each year. Lines depict the mean fit of the model. Points depict raw EVI values (n = 6-8 observations per year). Each shade of green corresponds with a different year; lighter greens represent earlier years and darker greens represent later years. The 50% normalized EVI (subsequently called 50% green-up) was calculated as the inflection point of each line. G24 tended to leaf out slightly earlier than ONA in each year, although not statistically significant.
