## Appendix S7 for "Spring ephemeral *Erythronium umbilicatum* may not be vulnerable to phenological mismatch with overstory trees"

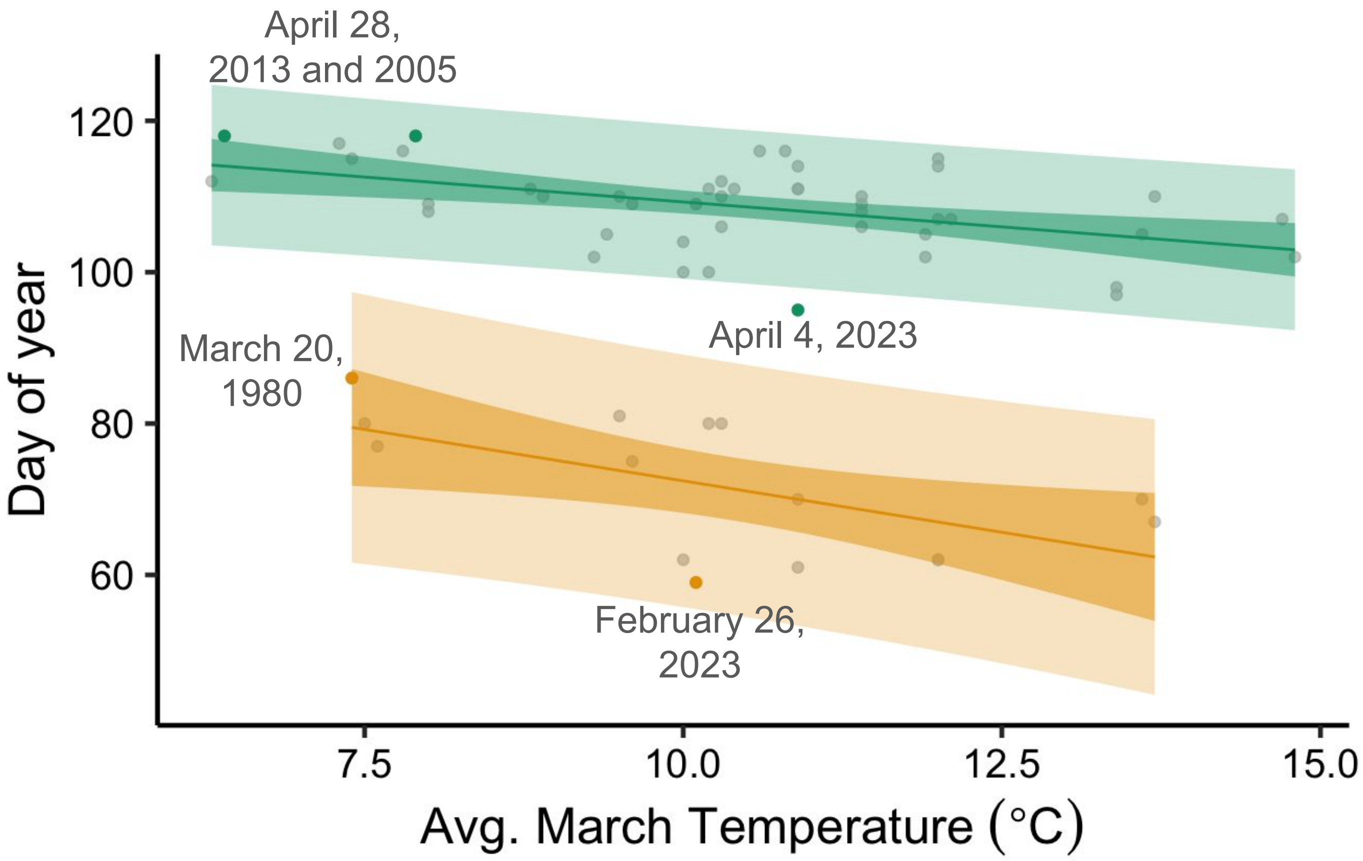


**Appendix S7.** Linear models to predict the day of year (DOY) that peak flowering and tree leaf-out will occur, given a mean March temperature. Gold line (lower) represents peak flowering trend and green line (upper) represents tree leaf-out. Points represent raw data, lines represent fitted model with 95% confidence (dark ribbon) and prediction intervals (light ribbon). Colored points and text highlight the minimum and maximum observed day of year that the phenological events occurred. All temperature data were collected from the PRISM database.
