## Appendix S8 for "Spring ephemeral *Erythronium umbilicatum* may not be vulnerable to phenological mismatch with overstory trees"

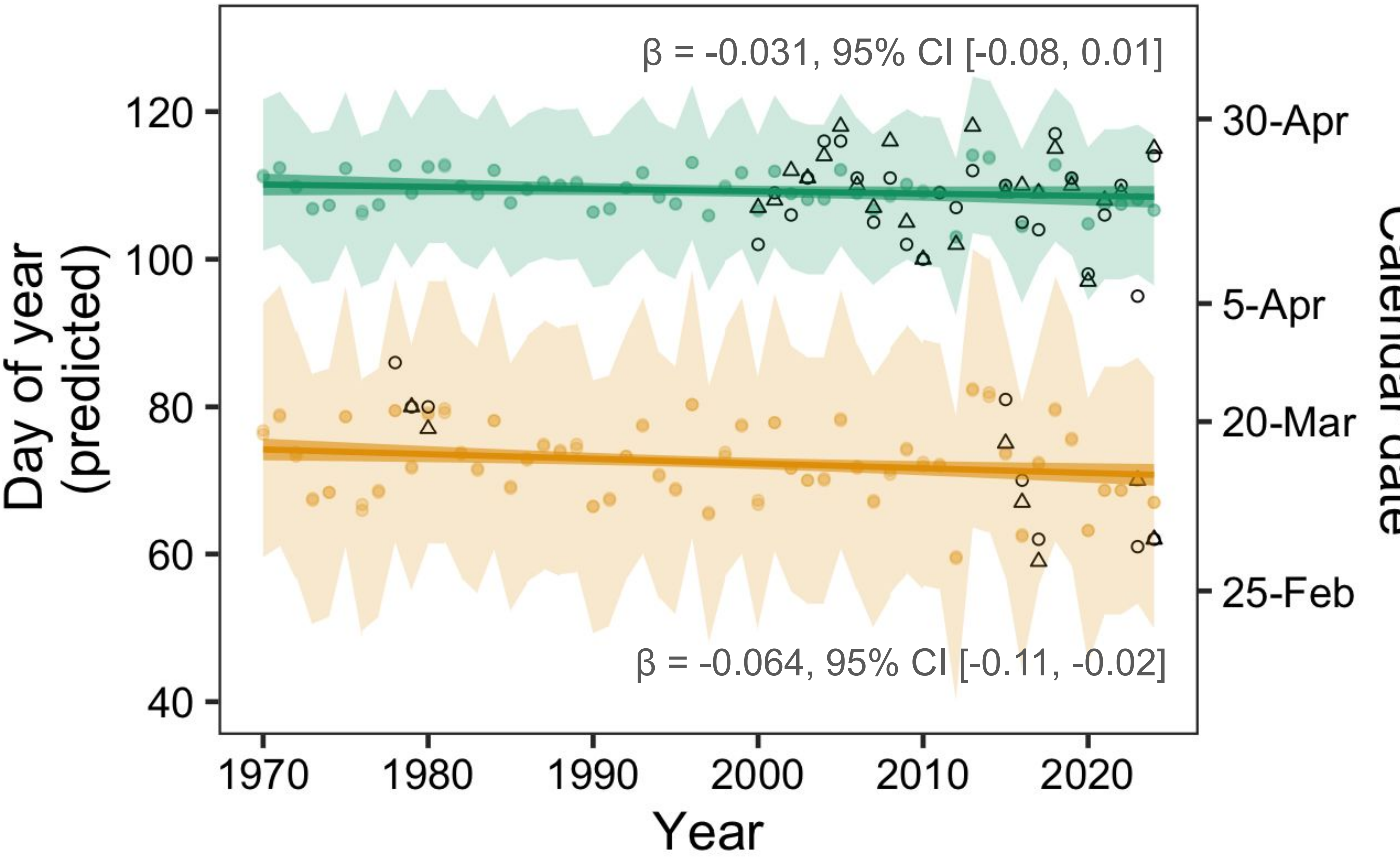


**Appendix S8.** Linear regressions were used to predict when a phenological event would occur given a mean March temperature for both flowering and tree leaf-out. The flowering phenology of *E. umbilicatum* (gold) but not the timing of tree leaf-out (green) is predicted to shift earlier in the year, although the slopes are not significantly different from each other ($F_{3, 216}$ = 1577, $R^2_{adj}$ = 0.96, P = 0.33). Colored points and jagged ribbons represent those predictions and 95% prediction intervals. Open circles (G24) and triangles (ONA) are the same raw data presented in Appendix S7, but were not involved directly in the linear fits shown here. Straight lines with 95% confidence intervals are outputs from linear model that correlates year with predicted day of year of phenological event.
