## Appendix S9 for "Spring ephemeral *Erythronium umbilicatum* may not be vulnerable to phenological mismatch with overstory trees"

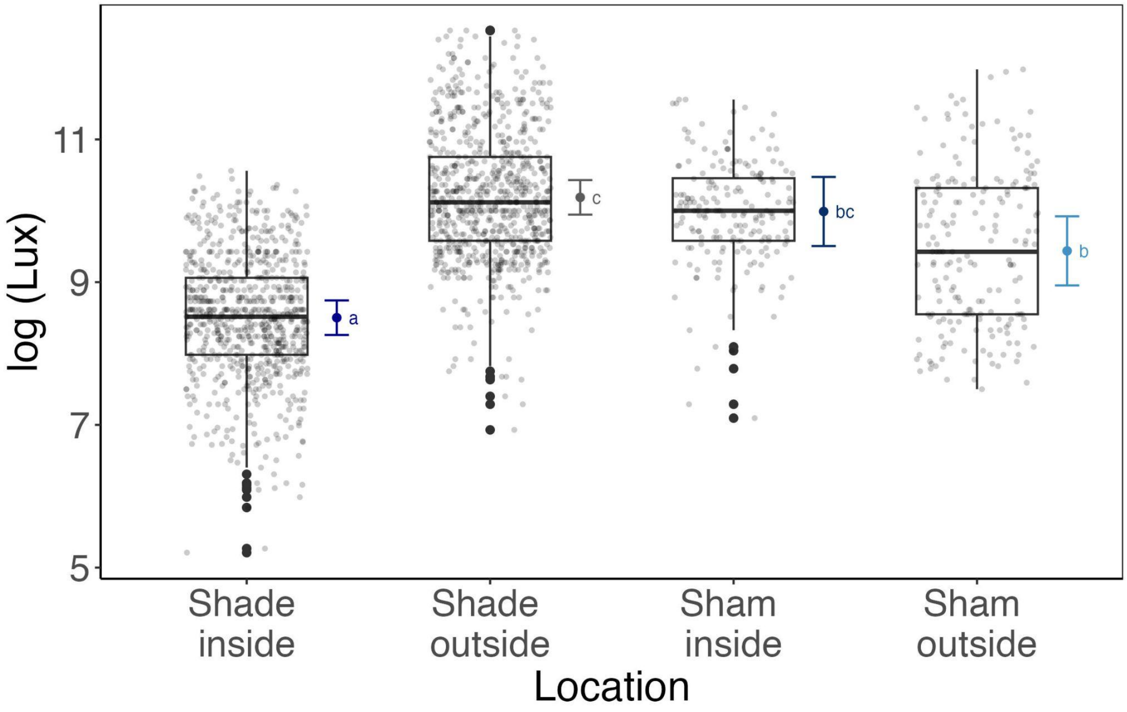


**Appendix S9.** The shade cloth significantly reduced the amount of light in the plots, while the sham light condition was not significantly different from ambient conditions ($\chi2 = 32.9$, df = 1, P = <0.0001). The light gray points depict the raw data on the log scale. Boxplots summarize the data, error bars represent the estimated marginal means and 95% CI.
