## Appendix S10 for "Spring ephemeral *Erythronium umbilicatum* may not be vulnerable to phenological mismatch with overstory trees"

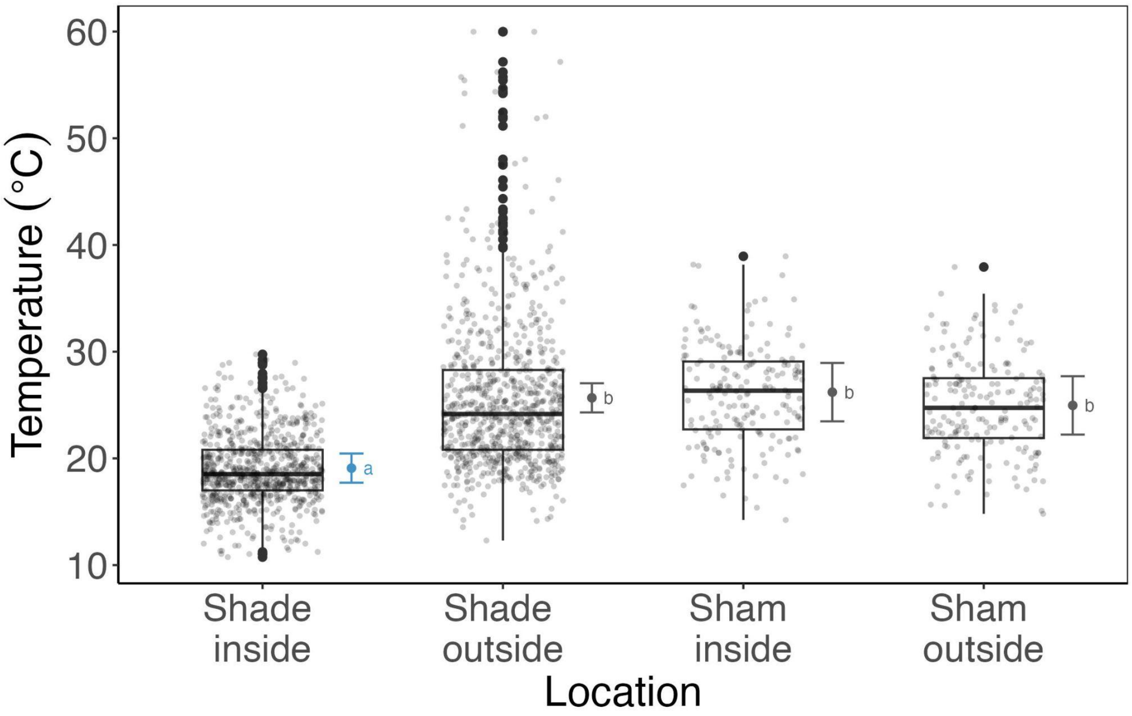


**Appendix S10.** The shade cloth significantly decreased daytime temperature compared to ambient and sham conditions ($\chi2 = 12.59$ df = 1,  P = <0.0001). Light gray points are data, boxplots summarize the data, error bars represent the estimated marginal means and 95% CI.
