## Appendix S11 for "Spring ephemeral *Erythronium umbilicatum* may not be vulnerable to phenological mismatch with overstory trees"

**Appendix S11.** The effects of observation day (center scaled), the number of days of extra shade, and the interaction between them in 2023 on the probability of senescence in 2023 and 2024. P-values significant at α = 0.05 are in bold font.

| Year | Predictor | Estimate | SE | z ratio | *p* | Prob at obs. shade days = 0 | Δ prob (shade days = 0 to 6) |
|---|---|---|---|---|---|---|---|
| 2023 | (Intercept) | -1.683 | 0.297 | -5.672 | **<0.001** | | |
| | Obs. DOY, center scaled | 3.034 | 0.302 | 10.041 | **<0.001** | | |
| | Shade days | -0.014 | 0.012 | -1.184 | 0.237 | | |
| | Obs. DOY * Shade days | 0.022 | 0.012 | 1.755 | 0.076 | | |
| 2024 | (Intercept) | 0.642 | 0.192 | 3.332 | **0.001** | | |
| | Obs. DOY, center scaled | 2.144 | 0.099 | 21.595 | **<0.001** | | |
| | Shade days | -0.017 | 0.008 | -2.249 | **0.025** | 0.47 | −5.5% |
| | Obs. DOY * Shade days | 0.005 | 0.004 | 1.184 | 0.236 | | |
