## Appendix S12 for "Spring ephemeral *Erythronium umbilicatum* may not be vulnerable to phenological mismatch with overstory trees"

**Appendix S12.** The effect of the number of days of extra shade in 2023 on the probability of flowering in 2024. P-values significant at $\alpha = 0.05$ are in bold font.

| **Predictor** | **Estimate** | **SE** | **z ratio** | ***p*** | **Prob at shade days = 0** | **Δ prob (shade days = 0 to 6)** |
|---|---|---|---|---|---|---|
| (Intercept) | -1.325 | 0.327 | -4.052 | **<0.001** | | |
| Shade days | -0.046 | 0.015 | -2.995 | **0.003** | 0.22 | −20.5% |
